# Structural Brain Correlates of Creative Personality

**DOI:** 10.1101/2021.01.06.425561

**Authors:** Divya Sadana, Rajnish K. Gupta, S. S. Kumaran, Sanjeev Jain, Jamuna Rajeswaran

**Author notes:** These authors contributed equally to this work (Divya Sadana, Rajnish K. Gupta). **Corresponding Author:** Dr. Jamuna Rajeswaran.

## Abstract

Creative individuals and their enigmatic personalities have always been a subject of fascination. The current study explored the neuroanatomical basis of creative personality using voxel-based morphometry. The sample comprised of two groups - Creative (CR) group (professional creative artists) and matched controls with no demonstrated artistic creativity (NC) with 20 participants in each group, in the age range of 20-40 years, right-handed, and had minimum average intelligence (IQ > 90). Participants in CR were selected using the creativity achievement questionnaire, creativity was assessed using the Wallach & Kogan test of creativity, and personality was administered using NEO-FFI. Results indicated significantly higher openness to new experiences in CR which positively correlates with the right middle frontal gyrus. An increased grey matter volume in the inferior frontal gyrus, anterior cingulate gyrus in CR, pointing towards the integration of cognitive and imaginative processes that might be implicated in creative personality.

## Introduction

Creativity is the quintessence of human life. However, the vastness and the versatility of this ability make it implausible to be explained by a single definition and more than 60 definitions of creativity exist in the psychological literature (Amabile, 1996; Runco, 2004; Taylor, 1988). The most consensual understanding of creativity delineates it as an interaction of personality and cognitive aspects of an individual leading to the production of an idea or a product that is both novel and useful (Gough, 1979; Pfeiffer, 2002; Rhodes, 1961; Runco, 2007). The nascent attempts in creativity research were aimed at unraveling the aptitudes and personalities of creative individuals.

Feist conducted the first meta-analytic review exploring the associations between creativity and personality emphasizing that while creativity is the study of a person’s unique ideas/ products generated from them, personality is the study of what makes a person unique. His findings indicated that creative people are more open to new experiences, less conscientious, more self-accepting, hostile, and impulsive (Feist, 1998). Since then, a plethora of research has emerged attempting to understand what aspects of personality are common across creative individuals and could play a vital role in nurturing the creative process. Openness to new experiences and Extraversion have been positively associated with creativity across numerous studies (Batey, Furnham, & Safiullina, 2010; Chamorro-Premuzic & Reichenbacher, 2008; Feist, 1998; Furnham, Crump, Batey, & Chamorro-Premuzic, 2009). Low Conscientiousness and high Neuroticism (Batey & Furnham, 2006; J. C. Kaufman, Cole, & Baer, 2009; Silvia, Kaufman, Reiter-Palmon, & Wigert, 2011) have also been associated with creativity, although less consistently. It has been hypothesized that individuals with high extraversion, openness, and low conscientiousness would provide more fluent, varied and unique responses to divergent thinking tests (Batey, Chamorro-Premuzic, & Furnham, 2009; Batey & Furnham, 2006; Hughes, Furnham, & Batey, 2013) and also rate themselves higher in self-rated creativity tests. Thus, the sample of these creativity studies and how creativity is assessed might have a determining effect on its association with certain personality traits.

To overcome this limitation, recent studies used improvised research designs involving professionally established creative artists. Fink and Woschnjak assessed personality correlates in three different groups of professional dancers (ballet, modern/contemporary, and jazz/musical) and found that modern/ contemporary dancers who are often required to freely improvise on stage, exhibited relatively high levels of verbal and figural creativity, followed by jazz/musical and finally by ballet dancers who largely perform rehearsed moves. On the personality dimension, modern/ contemporary dancers were found to have lower levels of conscientiousness than the groups of ballet and jazz/musical dancers. Also, modern/contemporary dancers showed higher scores (though not significant) on the personality dimension of openness to experiences than the other two groups (Fink & Woschnjak, 2011). Another study assessed the personality correlates of creativity in musicians (M. Benedek, Borovnjak, Neubauer, & Kruse-Weber, 2014) and found that folk musicians were more extraverted than classical and jazz musicians. Authors attributed this to the working style of folk musicians who commonly play at sociable events and have good interactions with the audience. Also, jazz and folk musicians scored high on openness to new experiences than classical musicians. While these functional studies provided important insights, findings varied with each study population, measures used, and the specific focus of brain activity being measured.

In such a scenario, structural scans offer a unique opportunity to view and measure all brain systems simultaneously, rather than a narrow focus on only those systems that are utilized for the performance of a particular task during a specific functional scan (C. G. DeYoung et al., 2010). The burgeoning realm of personality neuroscience has opened new avenues by linking mechanisms underlying each personality trait to specific brain regions associated with these functions. Neuroimaging studies (C. G. DeYoung et al., 2010; C. G. DeYoung, Peterson, & Higgins, 2005; C. G. DeYoung, Shamosh, Green, Braver, & Gray, 2009) have conceptualized Big Five factors in terms of their psychological attributes, localizing it to specific brain regions. Extraversion was found to be associated with the volume of the medial orbitofrontal cortex (involved in processing reward information); Neuroticism was associated with reduced volume in dorsomedial PFC and left medial temporal lobe including the posterior hippocampus, and with increased volume in the mid-cingulate gyrus (involved in a threat, punishment, and negative affect); Agreeableness was associated with reduced volume in the posterior left superior temporal sulcus and with increased volume in the posterior cingulate cortex (regions involved in social information processing); Conscientiousness was associated with volume in the lateral prefrontal cortex (involved in planning and the voluntary control of behavior); and Openness to experience has been associated with prefrontal cortex (regions involved in working memory, abstract reasoning and control of attention).

While these personality correlates have been localized in normal young adults, the structural basis of the creative personality remains elusive with very few studies in this area. In one such study, Li used voxel-based morphometry to identify the brain regions underlying individual differences in trait creativity and reported increased gray matter volume in the right posterior middle temporal gyrus (pMTG) in creative individuals associating it with semantic retrieval (Li et al., 2014). Authors also found that although openness to experience, extroversion, conscientiousness, and agreeableness, all contributed to trait creativity, only openness to experience mediated the association between the right pMTG volume and trait creativity. Openness to experience, one of the Big Five factor has been a subject of fundamental research in creativity and was found to predict creativity in a wide range of domains (e.g., arts, sciences, and humanities) (Feist, 1998) and levels of analysis (e.g., creative thinking styles, hobbies, and accomplishments) (Feist & Barron, 2003; King, Walker, & Broyles, 1996; Silvia et al., 2008).

As the field of creativity is expanding with information from fields of neuroscience and personality psychology, there is a need to fill lacunae in existing research on three major dimensions-1) Sample-Existing research on this construct has used either eminence as a “de facto criterion” of creativity which was problematic because it represented only an extreme subgroup of creative individuals or employed college students as the sample and administered creativity tests on them. 2) Behavioural Measure-Using a test that meets the central characteristics of producing novel and useful responses to items rather than self-ratings 3) Neuroimaging measure-It is difficult to interpret, or integrate across, imaging studies that use diverse creative cognition measures, most of unknown reliability and validity, and report activity in different brain regions of interest. The assessment of neuroanatomical aspects using structural imaging may be more efficacious for investigations of (stable) trait creativity than fMRI. Therefore, the current study attempts to explore the personality correlates associated with creativity and localize it to specific regions in the brain. In the present study, personality was assessed in a group of professionally established creative artists and we attempted to identify possible neural substrates of their personality using a voxel-based morphometric approach. It was hypothesized that creative artists would have significantly high scores on extraversion, neuroticism, openness to experience, and low scores on conscientiousness factors. The brain areas mediating these personality factors could possibly be in frontal and temporal regions of the brain.

## Materials and methods

A matched case-control design with a cross-sectional assessment was used for the study.

### Participants

The sample comprised of two groups-Professional Creative Artists (CR) and controls with no demonstrated artistic creativity (NC) with n=20 each. All participants were in the age range of 20-40 years, right-handed (screened using Edinburgh Handedness Inventory) and had minimum average intelligence (IQ > 90 on Raven’s Progressive Matrices). The groups were matched on age (mean age of CR was 30.2 ± 5.47 years and NC was 30.05 ± 5.34 years); gender (16 males and 4 females in each) and years of education (16.4 ± 0.75 in CR and 16.2 ± 0.83 in NC group). The CR group included established creative artists with Pro-c creativity (Pro-c level was established through the Creativity Achievement Questionnaire) and a minimum experience of two years in the creative field. NC group comprised of healthy adults from the community with no demonstrated artistic creativity and scores below the criteria for the Pro-c level. These participants were also tested on the Wallach & Kogan Test of Creativity (WKC) which has been standardized on the Indian population (Paramesh, 1972). The responses on various subtests were scored on three dimensions-Fluency (number of responses given to each item in each subtest), Originality (number of unique responses given to each item in each subtest), and Flexibility (number of categories represented by responses given to each item in each subtest). Individuals with any medical, psychiatric, or neurological disorders; with metal implants, pacemakers, or having a history of claustrophobia were excluded from the study. Written informed consent and socio-demographic details were obtained from all the participants and the study was approved by the Institute Ethics Committee.

### Measures

CR group was selected using Creativity Achievement Questionnaire (CAQ) (Shelley H Carson, Peterson, & Higgins, 2005). A self-report measure of creative achievement comprising of 96 items that assess achievement across 10 domains of creativity (visual arts, music, dance, creative writing, architectural design, humor, theatre and film, culinary arts, inventions, and scientific inquiry). A score of five and above on a single domain indicated the professional competence of the artist.

Personality was assessed using the NEO Five-Factor Inventory (NEO-FFI). The 60-item scale is a self-report version of the NEO-PI-R (Costa & McCrea, 1992) and assesses the five big domains of personality, namely Neuroticism, Extraversion, Openness to Experience, Agreeableness, and Conscientiousness.

### Magnetic Resonance Imaging (MRI) data acquisition

All participants underwent magnetic resonance imaging (MRI) scans on a 3 Tesla MRI scanner (Skyra, Siemens, Erlangen, Germany). During the scan, earbuds and foam pads were used to reduce scanner noise and participants’ head motion. Participants were instructed to relax but not to fall asleep, and minimize body movement during the data acquisition. T1-weighted images were obtained in a sagittal orientation by using the Magnetization Prepared by Rapid Gradient Echo (MPRAGE) sequence. The scanning parameters were: repetition time (TR) = 1600ms, echo time (TE) = 2ms, resolution = 256 x 256, slices = 176, flip angle = 9°, slice thickness = 0.9mm, and voxel size = 0.937 x 0.937 x 0.9mm.

### Voxel-Based Morphometry (VBM) Analysis

The structural data were processed and analyzed using statistical parametric mapping (SPM12; http://www.fil.ion.ucl.ac.uk/spm/software/spm12/) with Computational Anatomy Toolbox (CAT12) toolbox (http://dbm.neuro.uni-jena.de/cat/) running in MATLAB R2016a (www.mathworks.com). A standard VBM processing procedure was used as implemented in CAT12. All T1 images were segmented into gray matter (GM), white matter (WM), and cerebrospinal fluid (CSF) and normalized using the Diffeomorphic Anatomical Registration Through Exponentiated Lie algebra (DARTEL) template from 555 healthy control subjects in the IXI-database. The voxel size for the normalized image was 1.5 x 1.5 x 1.5 mm^3^. After preprocessing, a quality check was performed to assess the homogeneity of the GM tissues. Subsequently, all modulated and normalized GM images were smoothed with an 8 mm full width at half-maximum (FWHM) Gaussian kernel.

All the statistical analyses were performed using CAT12. To identify the GM volume differences in creative and control groups, an independent two-sample t-test was performed. Age and gender were used as covariates of no interest. The modulated GM images were corrected for non-linear warping, using global scaling for the Total Intracranial Volume (TIV). Therefore, proportional scaling was performed on the grand mean of scaling values. A significant threshold at p < 0.001 (uncorrected) was corrected at the cluster-level for multiple comparisons with an extent threshold of 1201 voxels. Anatomical labeling of significant clusters was performed using the xjView toolbox (https://www.alivelearn.net/xjview/). A binary mask from the resultant image was used as an inclusion mask for the following regression analyses.

Thereafter, to examine the association between the subject’s personality inventory scores (NEO-FFI) and GM volumes, we performed multiple regression analyses using the CAT12 toolbox, with age and gender as nuisance covariates, while global scaling for TIV. A significant threshold of p < 0.05 (FWE-corrected) was corrected at the voxel-level for multiple comparisons with an extent threshold of 10 voxels. Additionally, a partial correlation analysis between GM volume data and each NEO-FFI category scores were performed using SPSS v20.0 (https://www.ibm.com/analytics/spss-statistics-software).

## Results

Data were analyzed using SPSS v20.0 for windows. The Shapiro-Wilk test was used to test the normality of distribution (Shapiro & Wilk, 1965). The data, which were normally distributed, were analyzed using a two-sample independent t-test, while for the data which did not meet the assumptions of normality, a similar non-parametric Mann Whitney U-test was used.

### Comparison of Groups on Behavioural Measures

The age (years), years of education, scores on CAQ, WKC, and NEO-FFI measures of the sample with Mean ± SD of the participants in the two groups are presented in Table 1. A value of p < 0.05 was statistically significant.

**Table 1.**
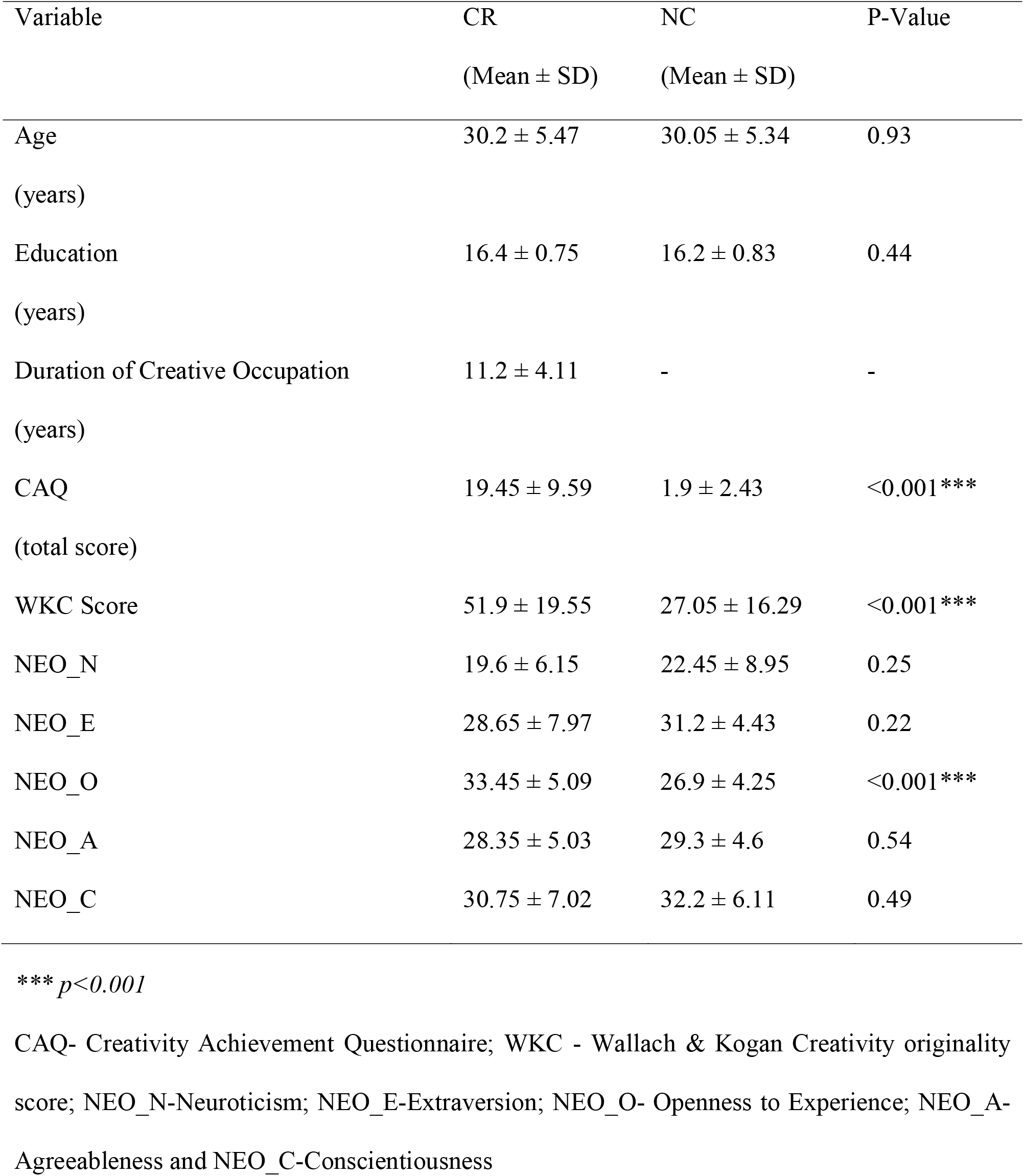
Comparison between Creative (CR; n=20) and Uncreative Controls (NC; n=20) on behavioural measures.

A statistically significant difference was found between the two groups on CAQ, WKC, and Openness to Experience aspects of personality between CR and NC. No statistically significant difference was found between the two groups on age, years of education, Neuroticism, Extraversion, Agreeableness, and Conscientiousness aspects of personality.

### Regional GM enhancement in the CR relative to NC

The CR group showed GM enhancement compared to NC in areas such as the inferior frontal gyrus (IFG) and anterior cingulate gyrus (Table 2, Figure 1). No significant enhancement was observed in GM in the NC group compared to the CR.

**Table 2.**
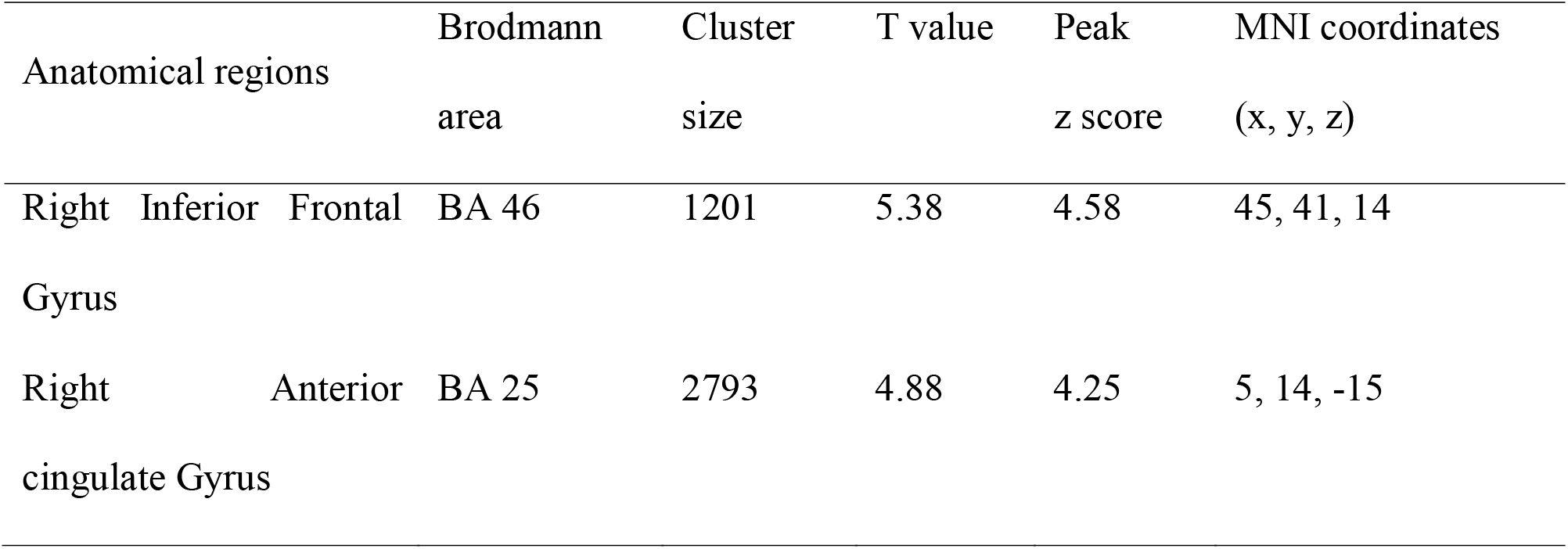
Anatomical regions with gray matter volume enhancement in the CR group compared with NC.

**Figure 1.**
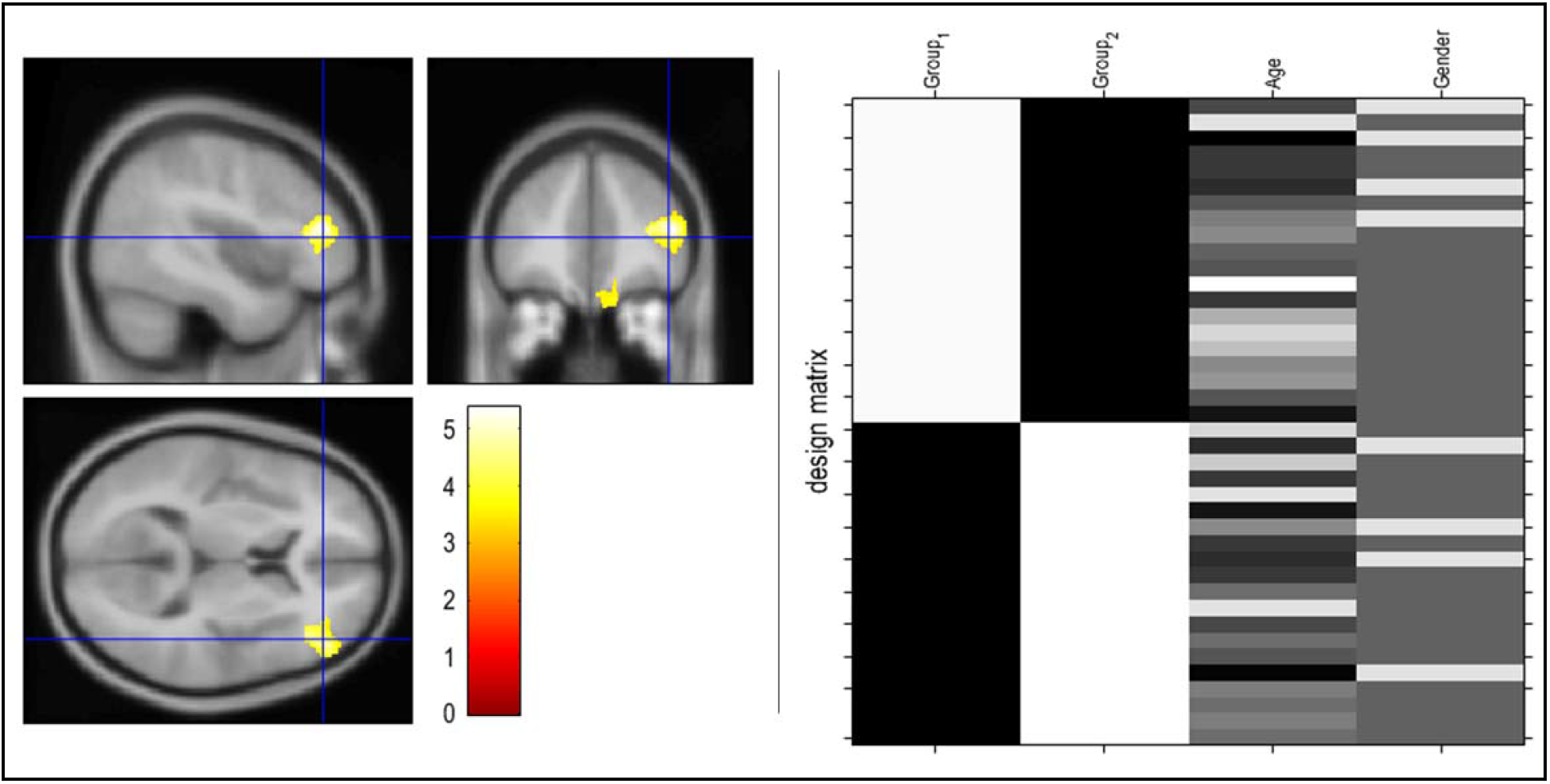
Group differences in GM volume. *Note*. Clusters reflects gray matter enhancement in the inferior frontal gyrus and Anterior Cingulate gyrus in the CR group compared with NC. The color bar represents the T-score: white indicates higher statistical significance than yellow or red. The threshold p < 0.001 was corrected for multiple comparison at cluster-level, extent threshold of 1201 contiguous voxels.

### Correlation analyses for GM volume and NEO-FFI scores

Multiple regression analyses found that NEO-O scores positively correlated with right middle frontal gyrus GM volume (peak coordinates: x, y, z = 42, 39, 20; cluster size = 11 voxels). None of the other NEO-FFI category scores significantly correlated with GM volume (Table 3, Figure 2). The scores for correlation analysis between GM volume and NEO-FFI are presented in Table 4.

**Table 3.**
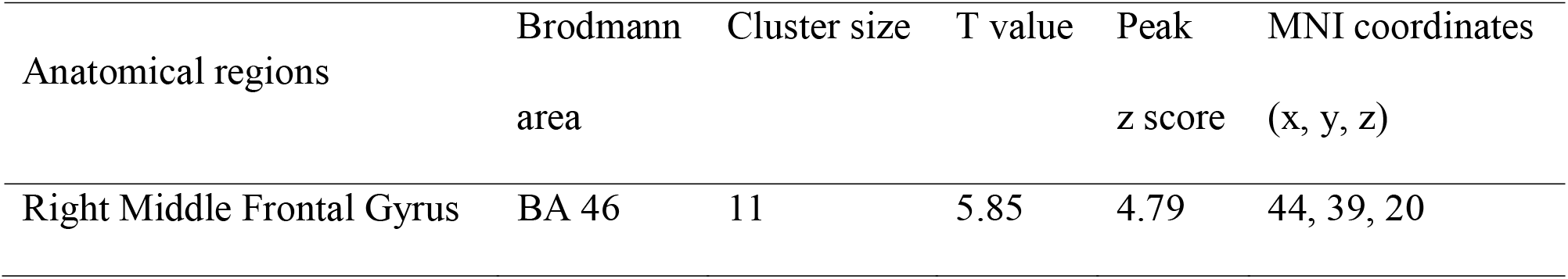
Anatomical regions with gray matter that significantly correlates with the NEO-FFI score.

**Figure 2.**
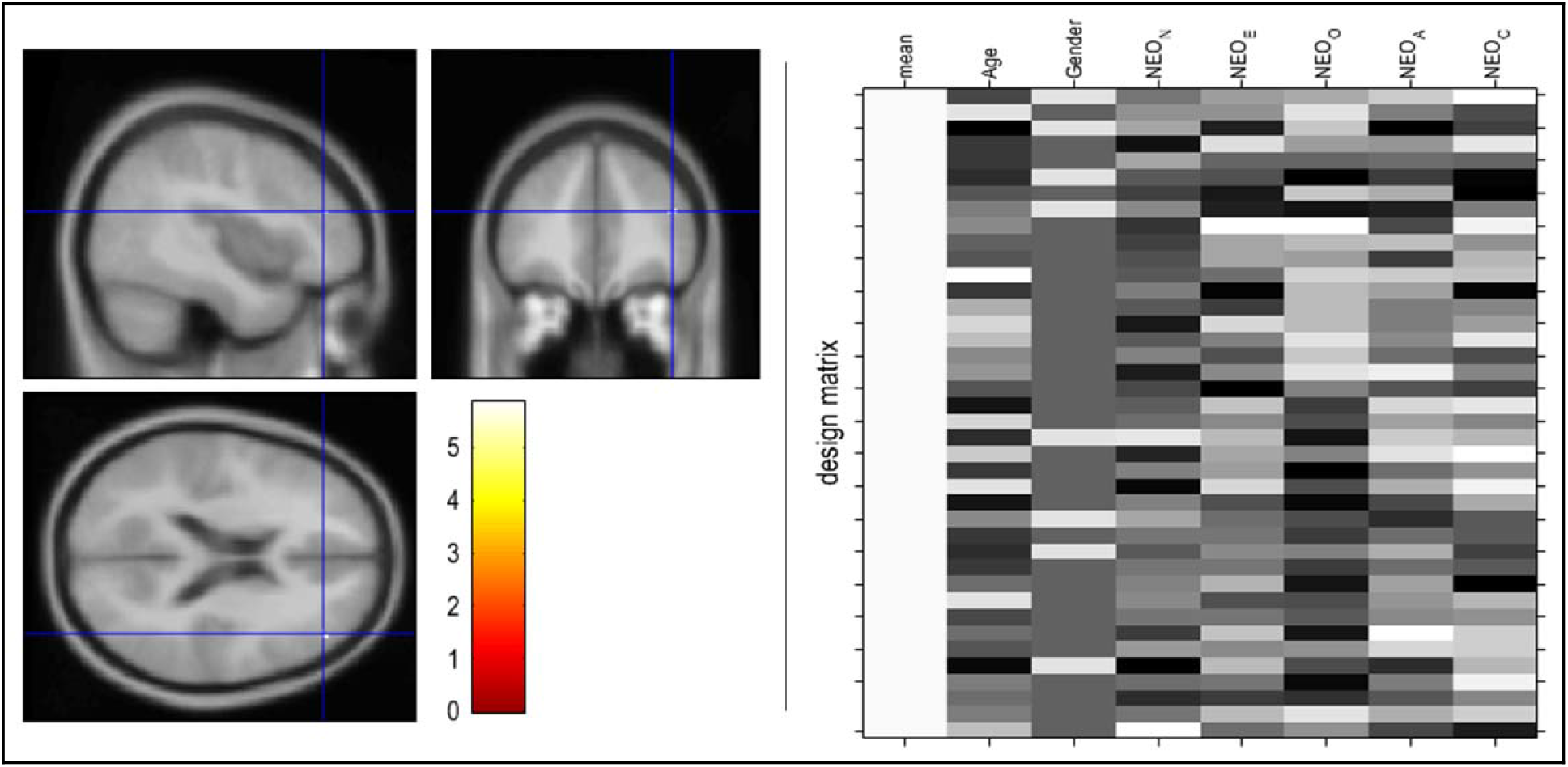
Multiple regression analysis using voxel-based morphometry to identify the brain region that significantly correlates with the NEO-FFI score. *Note*. The cluster right middle frontal gyrus (x=44, y=39, z=20) significantly correlates with NEO-O score. The color bar represents the T-score: white indicates higher statistical significance than yellow or red. The threshold p < 0.05 (FWE-correct) was corrected for multiple comparison at voxel-level.

**Table 4.** Correlation between Gray Matter Volume (GMV) and NEO-FFI score.

| Variable | Pearson Correlation Value |
| --- | --- |
| GMV and NEO-N | -0.287 |
| GMV and NEO-E | -0.045 |
| GMV and NEO-O | 0.324* |
| GMV and NEO-A | 0.103 |
| GMV and NEO-C | 0.004 |

\* Correlation is significant at the 0.05 level (2-tailed)

## Discussion

The study aimed at exploring personality factors associated with creativity and localizing them to specific anatomical regions in the brain. A matched case-control design was used, and creative professionals were compared with controls on big five factors of personality (NEO-FFI) and gray matter volume changes corresponding to these personality factors were identified.

The current study found significantly higher openness to experience in creative artists as compared to controls, corroborating the findings from existing literature (M. Benedek et al., 2014; Shelley H Carson et al., 2005; Dollinger, Urban, & James, 2004; Feist, 1998; Fink & Woschnjak, 2011; King et al., 1996; McCrae, 1987; Miller & Tal, 2007). Openness to experience has been consistently associated with creativity (Feist, 1998; Feist & Barron, 2003; King et al., 1996; Silvia et al., 2008). Deconstructing openness to experience as a trait, DeYoung, Quilty, and Peterson (2007) proposed that it has two primary aspects—openness (an imaginative, creative, and aesthetic aspect) and intellect (thinking and reasoning aspect) (Colin G DeYoung, Quilty, & Peterson, 2007). Openness/Intellect represents the capacity to process abstract and perceptual information flexibly and efficiently and includes four major dimensions namely cognitive ability, intellectual engagement, affective engagement, and aesthetic interest (S. B. Kaufman, 2013). Openness/Intellect has been consistently and positively associated with intelligence (C. G. DeYoung et al., 2005), a faculty that appears to be associated with the prefrontal cortex and with functions such as abstract reasoning, working memory and decision making (Gray, Chabris, & Braver, 2003). Thus, a higher degree of Openness/Intellect in creative artists suggests an increased tendency to expose themselves to diverse perceptual experiences and a larger capacity to assimilate these experiences intellectually and emotionally in their creative potential.

We analyzed the gray matter volume differences between the two groups and found that the CR group had higher GMV in right inferior frontal gyrus (BA46) and anterior cingulate gyrus (BA25). Our results corroborate findings from recent studies reporting significant positive correlations between creative potential and GMV in the inferior frontal gyrus (Zhu, Zhang, & Qiu, 2013), postulating its role in attentional processes (Zhang & Chiang-shan, 2012) and successful response inhibition (Aron, Robbins, & Poldrack, 2004) implicated in retrieval and selection of semantic concepts (Badre & Wagner, 2007; Blumenfeld & Ranganath, 2007). It is well known that divergent thinking is initially dominated by the retrieval of common, known ideas that are readily accessible whereas original and unique ideas occur at later stages in the ideation process (Beaty & Silvia, 2012; Mathias Benedek & Neubauer, 2013; Gilhooly, Fioratou, Anthony, & Wynn, 2007). IFG has been implicated in evaluating the originality of ideas by overcoming the readily accessible yet uncreative ideas and supporting the generation of new and more creative ideas (M. Benedek et al., 2014; Kleinmintz et al., 2018). Grabner, Fink, and Neubauer (2007) postulate that creative thinking is associated with enhanced anterior networks constituting lateral prefrontal, supplemental motor area, and cingulate gyrus and their research demonstrated an association between creativity and electrophysiology of the frontal and central regions of the right hemisphere (Grabner, Fink, & Neubauer, 2007). Cytoarchitecturally, BA46 roughly corresponds with the dorsolateral prefrontal cortex (DLPFC) and research has also indicated its involvement in functions like attention, verbal fluency, (Abrahams et al., 2003) working memory, and episodic long term retrieval (Ranganath, Johnson, & D’Esposito, 2003) and self-reflections in the decision-making process (Deppe, Schwindt, Kugel, Plassmann, & Kenning, 2005) thereby reiterating creativity as a higher-order cognitive function. Precentral gyrus is associated with the programming of motor movements and studies on creativity have consistently reported activations in this area during musical, visuospatial and mental imagery tasks of creativity (Berkowitz & Ansari, 2008; Boccia et al., 2015; Brown, Martinez, & Parsons, 2006; Milivojevic, Hamm, & Corballis, 2009; Pinho, de Manzano, Fransson, Eriksson, & Ullen, 2014).

In the final step using multiple regression analysis, we extrapolated all five personality factors and correlated them with gray matter volume changes in our participants. Results indicated that openness to experience was positively associated with increased regional GMV in the right middle frontal gyrus while none of the other NEO-FFI category scores significantly correlated with GM volume. Some studies in the area of personality neuroscience have found a negative correlation between prefrontal cortex volume and trait openness to experience (Li et al., 2014; Vartanian et al., 2018) suggesting thinking outside the box is enabled when the person’s ability to filter the contents of thought is reduced. However, other studies have considered openness to experience as the broadest domain, including a mix of traits relating to intellectual curiosity, intellectual interests, perceived intelligence, imagination, creativity, artistic and aesthetic interests, emotional and fantasy richness, and unconventionality (Batey & Furnham, 2006; S. H. Carson, Peterson, & Higgins, 2003; Feist & Barron, 2003; Hirsh, DeYoung, & Peterson, 2009; Silvia, Nusbaum, Berg, Martin, & O’Connor, 2009). Necka & Hlawacz (2013) compared bankers and artists in an effort to understand artistic temperament and exploring the relationships between creativity and temperament. Results indicated that the artists had a tendency toward activity, a “tendency to initiate numerous activities that lead to, or provoke, rich external stimulation (Nęcka & Hlawacz, 2013).” This vastness and richness of input create a corresponding richness of output, as individuals who “score high on activity tend to have many diverse experiences that may be used as a substrate for divergent thinking and creative activity.” This resonated with the responses to creativity data obtained in the present study. The CR group not just gave the most original responses, but the responses also exhibited an underlying diverse information base. For example, a CR participant reported “enso” as one of the responses to the test item asking for things that are round in shape. In Zen Buddhism, an ensō is a circle that is hand-drawn in one or two uninhibited brushstrokes to express a moment when the mind is free to let the body create. Now, this response is not merely an output of high intelligence, rather it also involves intellectual curiosity, a curiosity to inquire about phenomena in diverse cultures, and affective and aesthetic engagement. With a wide array of information available, against a backdrop of high analytic and reasoning ability, it becomes easier for CR individuals to provide unique, original responses on tests of divergent thinking. Neuroanatomical studies are based on the premise that repetitive regular performance of certain tasks produces structural changes in the areas mediating those tasks as a function of neuronal plasticity. CR group, comprising of professionally creative individuals could be temperamentally more inclined to explore diverse experiences and events, engaging intellectually and aesthetically, thereby having enhanced gray matter volume in regions mediating creativity.

Taken together, our results suggest that the basic personality trait of openness to experience exposes the individual to an array of experiences which when intelligently assimilated with contextual and emotional stimuli could lead to creative accomplishments. The gray matter volume enhancements in prefrontal regions and anterior cingulate gyrus point towards the integration of cognitive and imaginative processes that might be implicated in creative personality.

The study aimed at exploring the personality of professionally established creative artists and localizing them to specific regions in the brain. A matched case-control design, a sample of professionally creative artists and stringent statistical analyses lend methodological rigor and ecological validity to the findings of this study. However, a small sample size, few female participants and cross-sectional assessments are limitations of this study. Future larger cohort longitudinal studies could explore personality development, a manifestation of certain traits and its association with the refinement of the creative process.

## Funding

This study was supported by the Department of Science and Technology, Ministry of Science and Technology, India.

## Conflict of interest

None

## Notes

### Competing Interest Statement

The authors have declared no competing interest.

